# cCMP and cUMP stimulate the acid phosphatase activity of AphA in *Haemophilus influenzae*

**DOI:** 10.1101/2023.12.14.571460

**Authors:** Kristina Kronborg, Yong Everett Zhang

**Author notes:** Correspondence: Yong Everett Zhang.

## Abstract

We recently detailed the competitive inhibition of cyclic AMP (cAMP) on three periplasmic enzymes, AphA, NadN, and Hel, in *Haemophilus influenzae* Rd KW20. This inhibitory effect is vital for orchestrating the nutritional growth and competence development in KW20. Here, we extended the study to the three other second messengers, i.e., cyclic GMP, cyclic UMP, and cyclic CMP, each sharing structural similarities with cAMP. Notably, cGMP competitively inhibits AphA’s acid phosphatase activity akin to cAMP. In contrast, both cUMP and cCMP stimulate AphA’s phosphatase activity in a dose-dependent manner. This novel finding underscores the intriguing opposing effects of cyclic purine and pyrimidine nucleotides on AphA, suggesting potential intricate biological crosstalk among these second messengers.

## Introduction

Periplasmic non-specific acid phosphatases are enzymes that function as scavengers of various organic phosphomonoesters. Since phosphorylated compounds cannot cross the inner membrane of Gram-negative bacteria, these phosphatases allow the cell to take up extracellular compounds for use in various metabolic processes. One such phosphatase is AphA with homologs in both *E. coli, S. typhimurium* and *H. influenzae*. Recently, we found that the AphA homolog in both *E. coli* and *H. influenzae* is competitively inhibited by the well-known secondary messenger, cyclic AMP (cAMP) (1). In the same study, we reported that cyclic GMP (cGMP) seems to affect AphA activity as well; however, the molecular effect of cGMP on AphA was not studied further.

Furthermore, two additional cyclic nucleotides, namely the non-canonical second messengers cyclic CMP and cyclic UMP, have recently been acknowledged as authentic secondary messengers in bacteria (2). These molecules had long faced skepticism due to the absence of their dedicated synthetases (3). In bacteria, both cCMP and cUMP stimulate their specific effector proteins, mediating abortive phage infections. Given the similar chemical structures to cAMP, here we tested the effect of cGMP, cCMP, cUMP on the catalytic activity of *H. influenzae* AphA_Hi_.

## Results and Discussion

**Figure 1B** shows that cGMP appears to competitively inhibit AphA_Hi_, similar to the effects of cAMP (**Figure 1A**). Conversely, both cCMP and cUMP exhibit a dose-dependent stimulation of AphA_Hi_ phosphatase activity (**Figure 1C,1D**). This intriguing discovery underscores the opposing effects of purine and pyrimidine cyclic nucleotides on AphA_Hi_ activity. This new discovery indicates potential interesting biological circumstances. The recent identification of specific cC/UMP cyclases (Pyrimidine nucleotide cyclase, Pyc) in Bacteria and their pivotal roles in antiphage defense, is indeed a milestone in the field (2). However, it is noteworthy that these Pyc enzymes have a relatively narrow phylogenetic distribution, as indicated by the authors (2). Additionally, many extensively studied model (pathogenic) bacteria, such as *E. coli, H. influenzae*, and *Vibrio cholerae*, do not possess the Pyc enzymes. Furthermore, it is important to consider that while host mammalian cells may produce cC/UMP (possibly through the promiscuous activities of cA/GMP cyclases) and excrete them, their concentrations are anticipated to be lower than that of cA/GMP.

**Figure 1.**
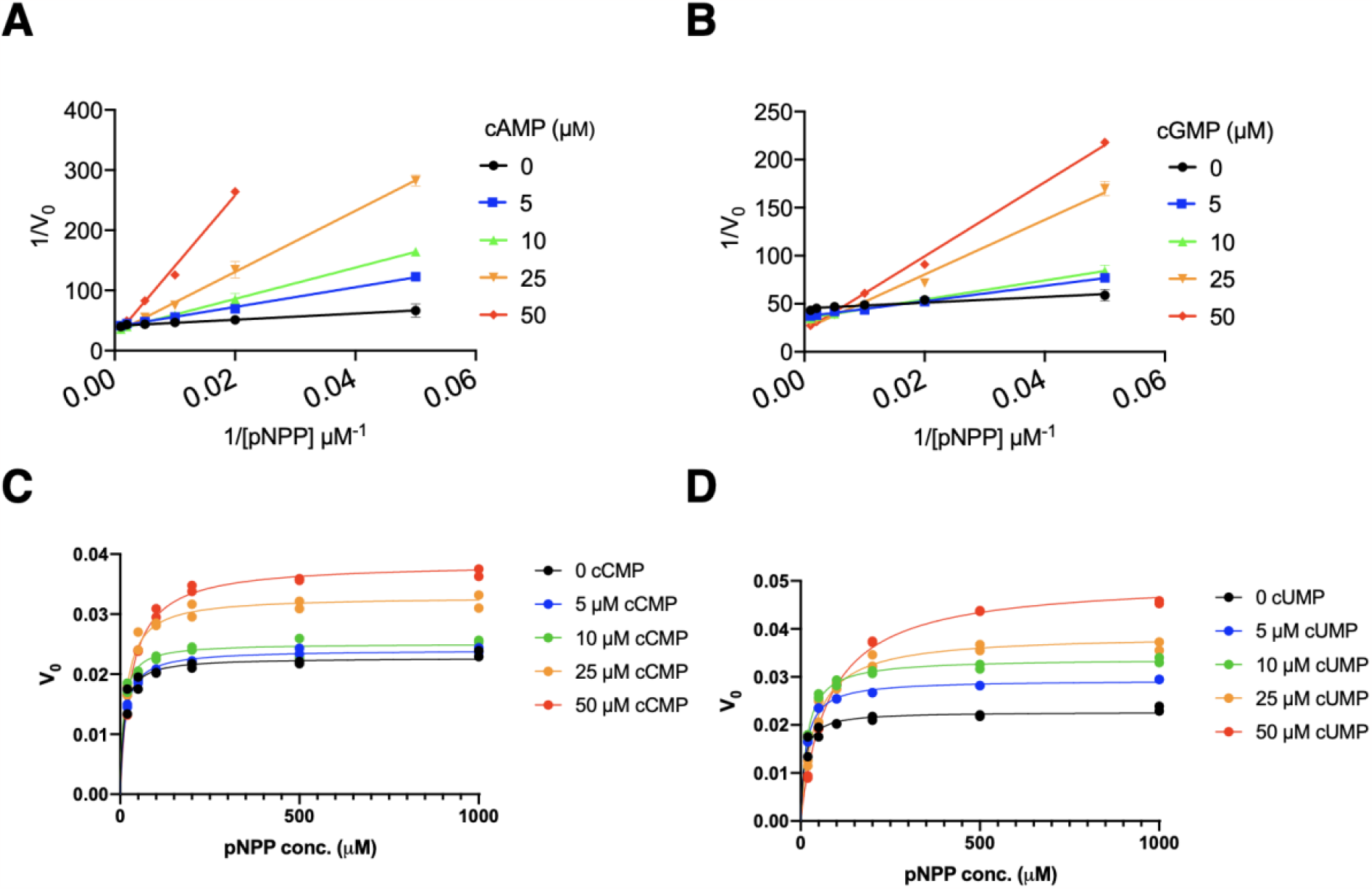
AphA activity is stimulated by cCMP and cUMP and inhibited by cAMP and cGMP. (**A**,**B**) Lineweaver-Burk plot of *H. influenzae* AphA kinetics in the presence of varied concentration of cAMP (**A**) and cGMP (**B**). (**C**,**D**) Michaelis-Menten plots of *H. influenzae* AphA in the absence and presence of cCMP (**C**), and cUMP (**D**). pNPP was used as substrate and formation of the yellow p-nitrophenol was quantified spectrophotometrically at 405 nm. Data represent two biological replicates.

On the other hand, there is the intriguing possibility that extracellular cC/UMP may stimulate the activities of AphA_Hi_ (as well as NadN and Hel), facilitating the nutritional uptake of NAD and nucleotides, ultimately impacting the growth of *H. influenzae*. This opens the possibility that the dynamic regulation of these three enzymes by various cyclic nucleotide molecules produced by KW20, other microbiota, and host cells collectively influences when and how KW20 becomes competent to uptake external DNA. This broader perspective on inter-species communication is potentially relevant and warrants further exploration. At last, but not least, the mechanistic details of cU/CMP’s stimulatory effect on AphA_Hi_ requires further dedicated study.

## Experimental procedures

*Protein purification and enzymatic assays were performed as (1)*.

### Preparation of chemicals

cAMP (SIGMA, A9501) was dissolved in Milli Q (MQ) H_2_O. cUMP (BioLog, 56632-58-7) and cCMP (BioLog, 54925-33-6) were each dissolved in a buffer of 25 mM Tris-HCl (pH 7.4) and 100 mM NaCl to a concentration of 250 mM and were then diluted to the desired concentrations in MQ H_2_O. p-nitrophenyl phosphate, pNPP, (SIGMA, S0942) was dissolved in 50 mM NaOAc to a concentration of 50 mM and was then serially diluted in the same buffer to the desired concentrations.

## Acknowledgement

This study was supported by a Danmarks Frie Forskningsfond (2032-00030B) to Y.E.Z.

---

**Table 1:** Bacterial strains used in this study.

| Strain | Relevant features | Reference |
| --- | --- | --- |
| YZ254 | BI21 DE3, pET28a-his.aphAhi, kanamycin | (1) |

## Notes

### Competing Interest Statement

The authors have declared no competing interest.

